# A small single-domain protein folds through the same pathway on- and off- the ribosome

**DOI:** 10.1101/347864

**Authors:** Emily J. Guinn, Pengfei Tian, Mia Shin, Robert B. Best, Susan Marqusee

**Author notes:** Correspondence should be addressed to S.M.

## Abstract

*In vivo*, proteins fold and function in a complex environment where they are subject to many stresses that can modulate protein energy landscapes. One aspect of the environment pertinent to protein folding is the ribosome, since proteins have the opportunity to fold while still bound to the ribosome during translation. We use a combination of force and chemical denaturant (chemo-mechanical unfolding), as well as point mutations, to characterize the folding mechanism of the src SH3 domain both as a stalled ribosome nascent chain and free in solution. Our results indicate that src SH3 folds through the same pathway on and off the ribosome. Molecular simulations also indicate that the ribosome does not affect the folding pathway for this small protein. Taken together, we conclude that the ribosome does not alter the folding mechanism of this small protein, which appears to fold at the mouth of the ribosome as the protein emerges from the exit tunnel. These results, if general, suggest the ribosome may exert a bigger influence on the folding of multi-domain proteins or protein domains that can partially fold before the entire domain sequence is outside the ribosome exit tunnel.

## Introduction

Proteins function in a complex cellular context where they are exposed to countless perturbing conditions that influence the folding process (*1-3*). Variations in environmental conditions such as temperature, solutes and mechanical stress are well known to modulate protein energy landscapes (*4-10*). In order to fully understand protein folding *in vivo*, it is essential to consider all conditions that a protein may experience in the cell, from synthesis to degradation.

A fundamental influence on the folding of every protein is the ribosome, which synthesizes proteins via translation of an mRNA transcript. Because folding often occurs on a faster timescale than translation (*11*), proteins have the opportunity to fold as they are being translated. Yet, we are just beginning to learn how the ribosome affects the folding process (*11-18*). There are many aspects of folding that may differ between a free protein and a protein tethered to the ribosome during translation. For instance, the emerging peptide chain can explore conformational space before the entire protein is synthesized. Indeed, small protein domains have been shown to fold in the mouth of the ribosome or even deeper in the ribosome exit tunnel, creating a strong spatial and steric constraint on the folding process (*18-20*). Because the ribosome is a highly-charged macromolecule, short-range chemical interactions, such as hydrophobic interactions and hydrogen bonding, and longer range coulombic interactions of the emerging protein with the ribosome could also affect the folding process (*21*).

Recent advances in experimental methodologies have begun to shed light on how the ribosome alters protein folding. Structural studies on ribosome nascent chain complexes (RNCs) using NMR and cryoEM have shown that, while secondary structure and simple tertiary motifs can form inside the ribosome exit tunnel, most proteins cannot find their native structure until they are outside the tunnel (*12, 18, 20*). Recent experiments harnessing the ability of the force generated by folding to release stalled nascent chains provide insight into the point during translation when different types of proteins fold (*17-19, 22*). Pulse proteolysis and optical trapping experiments have been used to explore the energy landscape of RNCs outside the exit tunnel, showing that the ribosome significantly alters the stability and dynamics of proteins and that the magnitude of this effect depends on the distance from the ribosome exit tunnel (*13, 14*).

Moreover, translation rates, which can be modulated by codon usage and mRNA secondary structure, affect the folding efficiency of a protein, in part by altering the time during which the protein interacts with the ribosome (*23-27*).

How does the ribosome affect the protein-folding pathway? For multi-domain proteins, the vectorial nature of co-translational folding can affect the pathway by altering which domain folds first (*15, 28, 29*). However, little work has been done to explore how the ribosome affects the actual folding pathway, or folding barrier, of a single protein domain. It is important to understand how this pathway differs for folding on- and off- the ribosome - the ribosome could favor or disfavor pathways prone to misfolding or aggregation and different residues may contribute to the rate of folding and unfolding under these different conditions. Folding simulations of an Ig domain on- the ribosome compared to experimental data off- the ribosome suggest that the ribosome does not affect its folding pathway (*19*). However, to date there are no experimental data probing the folding barrier (i.e. the transition state) of a single protein domain on the ribosome.

Previously, we combined chemical denaturant and force in a technique termed chemo-mechanical unfolding, as a means to probe the folding pathway of the src SH3 domain (*5, 6*). The src SH3 domain can access at least three different folding routes and small changes in chemical denaturant, force and sequence can alter the flux between these paths. We now use this same chemo-mechanical unfolding approach to explore another environmental condition: the ribosome. Employing the methodology developed by Kaiser et al (*13, 30*), we use optical tweezers to apply force on stalled RNC complexes containing the src SH3 domain. To distinguish between the different folding pathways of src SH3 we measure four parameters that characterize the transition state: urea m^‡^-values, force x^‡^-values, mutational ϕ-values, and the extrapolated unfolding rate in the absence of force or denaturant. We measure those same parameters for the same construct on and off the ribosome (RNCs and free protein) and use them to determine which pathway src SH3 folds through when it is tethered to the ribosome. Our results indicate that this src SH3 construct folds through the same pathway on and off the ribosome. Simulations of src SH3 folding both on and off the ribosome support these findings. This indicates that small single domain proteins can access the same folding pathway in both co-translational and refolding conditions and so perhaps the ribosome more easily modulates the folding pathway of larger multidomain proteins. This hypothesis is also supported by simulated ϕ values for the co-translational folding pathway of the small Ig domain, titin I27 (*19*).

## Results and Discussion

### Protein constructs for optical trap studies on- and off- the ribosome

Protein constructs used in the optical tweezers are tethered between two beads via DNA handles attached to genetically-encoded Avi and ybbr peptide tags (*31, 32*); one bead is held in a micropipette and the other is held in an optical trap to apply force (Figure 1a). To monitor free src SH3, these tags are encoded at the N- and C-termini of the protein. To monitor RNCs, the Avi tag is encoded in the N-terminus of src SH3 and the ybbr tag is encoded on protein L17 of the ribosome (the C-terminus of the RNC construct does not contain a peptide tag, ensuring that our optical trapping experiments will not capture any free protein released by the ribosome) (*13*). The RNC constructs (Figure 1b) contain a strong variant of the SecM stalling sequence (*33*) at the C-terminus to tether the protein to the ribosome and a Glycine-Serine (GS) linker to ensure that src SH3 is fully outside the ribosome exit tunnel. To alter the distance between src SH3 and the ribosome, we use constructs with either 10 or 20 GS repeats which, together with the 18 amino acid SecM sequence, create linkers of 38 and 58 amino acids respectively between src SH3 and the peptidyl transferase center (these constructs are referred to as SH3(38) and SH3(58)). Previous work has shown that a 38 amino acid linker should be sufficient to ensure that the protein sequence is entirely outside of the ribosome exit tunnel. Effective cleavage was seen in an RNC construct with a protease cleavage site N-terminal to the linker, supporting the expectation that the protein is not buried and inaccessible inside the exit tunnel (*14*). These GS linkers are also added to the free protein construct (Figure 1b) to make the free protein and RNC constructs as similar as possible.

**Figure 1.**
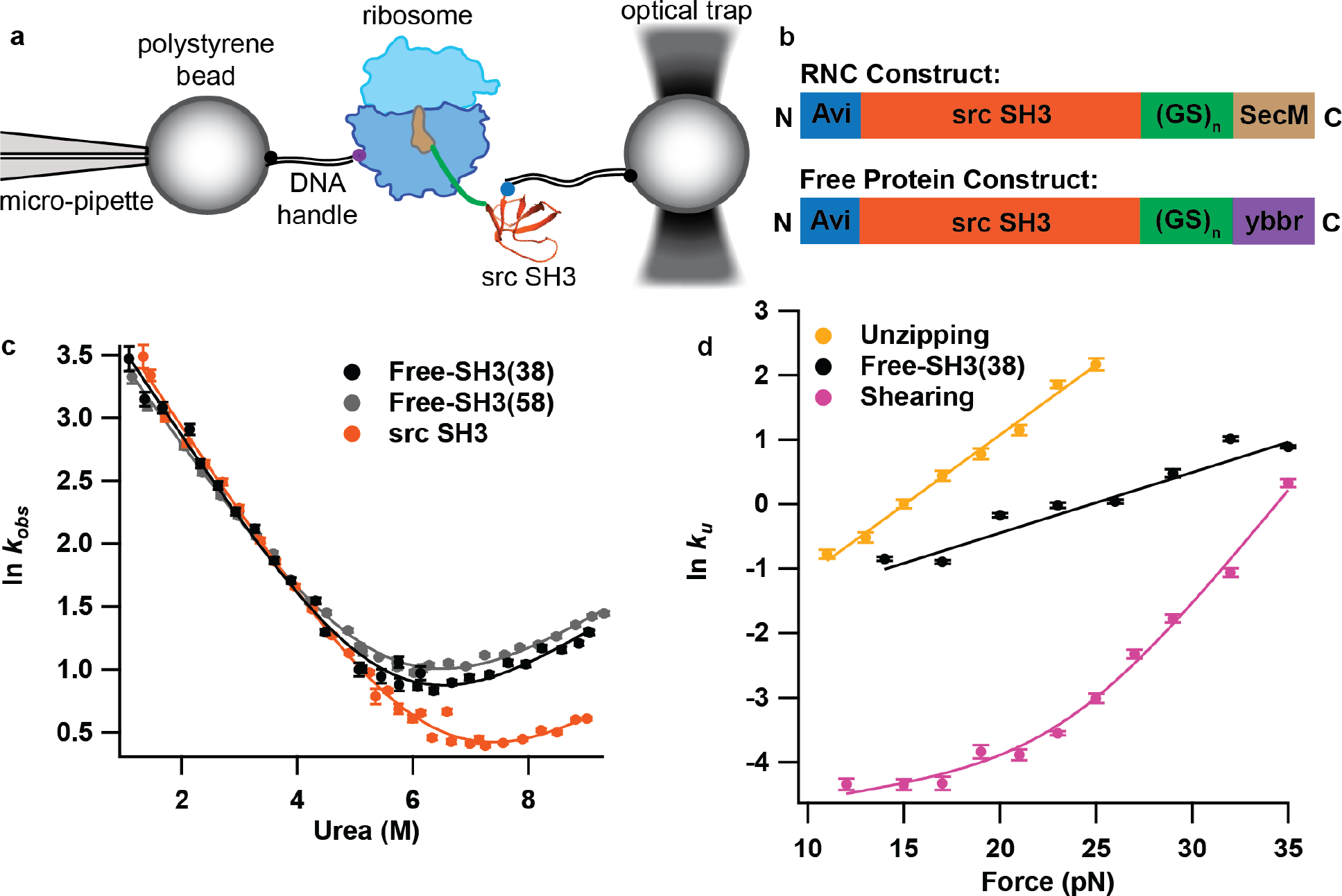
Overview of experimental setup and src SH3 constructs studied. (A) Schematic diagram of the experimental setup used to apply force to stalled ribosome nascent chains (RNCs) using the optical tweezers. DNA handles are attached to the RNC via an N-terminal Avi tag and a ybbr tag on protein L17 of the ribosome. (B) Constructs used in these experiments for RNCs (top) and free protein (bottom). SH3(38) constructs contain a linker with 10 GS repeats and SH3(58) constructs contain a linker with 20 GS repeats. (C) Kinetic chevron plots of the natural log of observed relaxation rate against urea concentration for src SH3 without the tags and linkers used in this work, and for the free-SH3(38) and free-SH3(58) constructs. Data were collected in 25 mM Hepes 150 mM KCl 5mM Mg-Acetate pH 7.4 (HKM) buffer. (D) Plots of the natural log of the unfolding rate against force for free-SH3(38) compared to previously published data pulling in unzipping and shearing geometries. The unzipping and shearing data were collected in 100 mM Tris 250 mM NaCl pH 7 buffer. The free SH3_38aalinker_ data were collected HKM buffer.

### Ensemble based folding studies on free-SH3 constructs with linkers and tags

To determine if the added linkers and tags affect the folding of src SH3, we measured folding and unfolding kinetics for free-SH3(38), free-SH3(58) and src SH3 without the tags and linkers. Kinetic data were collected using stopped-flow fluorescence spectroscopy and were well fit by a single exponential for all of the constructs. The resulting chevron plots (In *k*_*obs*_ vs [urea]) (Figure 1c) were fit assuming a two-state model:

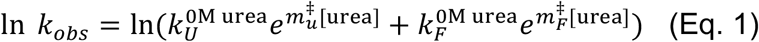

where *k*_*obs*_ is the observed relaxation rate, *k*_*U*_^0M urea^ and *k*_*F*_^0M urea^ are the unfolding and folding rates in the absence of urea and *m*_*U*_^‡^ and *m*_*F*_^‡^ are the unfolding and folding m^‡^-values (the slope of the natural logarithm of the unfolding or folding rate against urea concentration). While the tags and linkers increase the unfolding rate of SH3, they do not affect the folding rate, *m*_*U*_^‡^ or *m*_*F*_^‡^ (Table 1, Ensemble Data). Kinetic *m*^‡^-values are related to the change in accessible surface area for folding and unfolding to the transition state (*34, 35*). Since they are the same for all these constructs, the structure of the transition state is likely not altered by the tags and linkers. Therefore, the tags and linkers appear to increase the unfolding rate of src SH3 mostly by destabilizing the folded state. The SH3(58) unfolding rate is within error of the SH3(38) rate, suggesting that this change does not depend on the length of the linker.

**Table 1.** Results from kinetic analysis of ensemble and single molecule force experiments

| | Linker Length<br>(amino acids) | Urea $m_u^\ddagger$ ($M^{-1}$) | $x_U^\ddagger$ (nm) | $\ln k_U^{0pN, 0M\ urea}$ |
| --- | --- | --- | --- | --- |
| **Ensemble Data** |  |  |  |  |
| SH3<sup>a</sup> | 0 | $0.29 \pm 0.04$ | NA | $-2.03 \pm 0.38$ |
| Free-SH3<sup>b</sup> | 38 | $0.30 \pm 0.04$ | NA | $-1.47 \pm 0.31$ |
| Free-SH3<sup>b</sup> | 58 | $0.29 \pm 0.02$ | NA | $-1.23 \pm 0.15$ |
| Free-SH3 L24A<sup>b</sup> | 38 | $0.28 \pm 0.02$ | NA | $1.08 \pm 0.13$ |
| **Single Molecule Force Data** |  |  |  |  |
| Free-SH3<sup>b</sup> | 38 | $0.31 \pm 0.23$ | $0.37 \pm 0.02^d$ | $-2.12 \pm 0.16$ |
| RNC-SH3<sup>c</sup> | 38 | $0.33 \pm 0.23$ | $0.37 \pm 0.02^d$ | $-2.63 \pm 0.16$ |
| RNC-SH3<sup>c</sup> | 58 | NA | $0.37 \pm 0.02^d$ | $-2.76 \pm 0.19$ |
| RNC-SH3 L24A<sup>c</sup> | 38 | NA | $0.37 \pm 0.02^d$ | $-1.85 \pm 0.18$ |

a. Contains no tags of linkers.
b. Contains N-terminal Avi tag and C-terminal ybbr tag.
c. Contains N-terminal Avi tag and C-terminal SecM sequence.
d. These values were linked in global analysis.

### Effect of force on unfolding rates off the ribosome

Our approach requires that the DNA handles for optical trap studies are attached via genetically encoded peptide tags, imposing the restriction that the force is applied between the N- and C-termini of the protein. This differs from our previous work with src SH3, where we attached DNA handles to cysteines within the protein to pull along different geometries (shearing (A7C/N59C) and unzipping (R19C/N59C)) (*5*). To compare this new N- to C- pulling geometry to the previous shearing and unzipping geometries, we pulled on the free-SH3(38) construct and determined the unfolding rate as a function of force using force-jump experiments (*7*). The natural logarithm of the resulting unfolding rates are plotted against force and fit using the Bell equation (*36*):

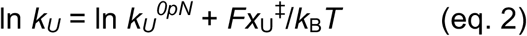

where *x*_U_^‡^ gives the effect of force on the unfolding rate, *k*_*U*_^*0pN*^ is the extrapolated unfolding rate in the absence of force, *k*_B_ is the Boltzmann constant and *T* is the absolute temperature. The *x*_U_^‡^ value is related to the distance between the native state and unfolding transition state along the mechanical reaction coordinate (*37*). Figure 1d compares this free-SH3(38) data to the previously published unzipping and shearing data. Unlike the upward curvature in the force dependence of ln *k*_*u*_ with a shearing force, which suggests parallel unfolding pathways (*5*), both the unzipping and free SH3(38) data can be approximated by a linear relationship between ln *k*_*U*_ and force, indicating that when pulled in these geometries, SH3 unfolds through a single pathway over the full range of force probed. The x_U_^‡^ and ln *k*_U_^0pN^ values differ between the unzipping and free-SH3(38) data; however, these data sets use different pulling geometries and buffers (in this work, we use a buffer compatible with RNCs), so we cannot determine if these differences are due to the different experimental conditions or represent different transition states. We showed previously that the unzipping pathway is likely the same as the pathway seen in ensemble experiments. Therefore, a comparison of the denaturant dependence of free-SH3(38) force data to ensemble kinetic data will help us to determine if this is the same as the unzipping pathway (see chemo-mechanical unfolding section below).

### Determining the effect of the ribosome on the force-dependence of unfolding

To explore the effect of the ribosome on folding, we apply force to RNC-SH3(38) and again use force-jump experiments to determine the unfolding rate as a function of force (Figure 2a) and fit to Bell’s model (eq 2). The force-dependent RNC-SH3(38) data run parallel to that for free-SH3(38), indicating that the x_U_^‡^ value is the same on- and off- the ribosome. Therefore, the ribosome does not alter the distance to the transition state, suggesting that src SH3 folds through the same pathway on and off the ribosome. The rates for the RNC data are slightly lower than the rates for free protein, indicating that the ribosome may affect the stability of the folded state or the transition state, as we discuss further below.

**Figure 2.**
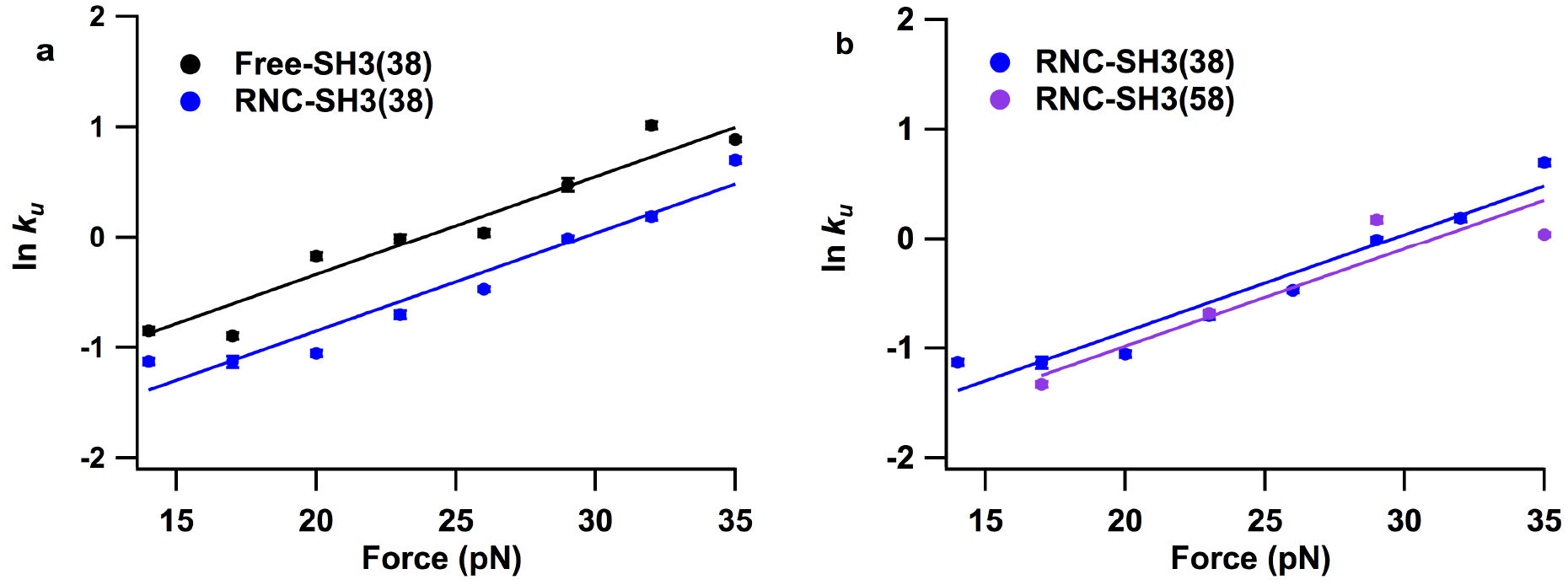
Effect of force of free-SH3 and RNC-SH3 constructs. Plots of the natural log of the unfolding rate against force for (A) free protein and RNCs with a 38 amino acid linker and for (B) RNCs with a 38 amino linker and a 58 amino acid linker.

To explore how increasing the distance from the ribosome affects the unfolding rate, we performed the same force-jump experiments with RNC-SH3(58). Plots of the RNC-SH3(38) and RNC-SH3(58) data (Figure 2b) look quite similar, suggesting that increasing the distance from the ribosome does not significantly affect the unfolding rate or pathway. Simulations (see below) suggest that our linker length of 38 amino acids represents the lower limit that allows folding of a RNC. Consequently, all subsequent experimental analysis will use only SH3(38) constructs.

### Using chemo-mechanical unfolding to compare the folding pathway on- and off- the ribosome

The above ensemble and force-dependent results (Figures 1 and 2) suggest that src SH3 may unfold through the same pathway on- and off- the ribosome and at different distances from the ribosome. To test this hypothesis directly, we turn to chemo-mechanical unfolding to determine an additional parameter, the *m*^‡^-value, that is also related to the structure of the transition state and so can help distinguish between pathways (*5, 34, 35*). Moreover, because *m*^‡^-values can be measured in standard ensemble experiments (Figure 1), they provide a means to compare the pathways seen in single molecule force data to the pathways seen in ensemble data.

In chemo-mechanical unfolding experiments, we simultaneously determine the effect of force and urea on unfolding of free-SH3(38) and RNC-SH3(38). We determine the unfolding rate as a function of force in buffer containing 1M urea to compare to the data collected in 0M urea. Figure 3 shows that urea affects the free protein and RNCs similarly; for both, the 1M urea data show a linear relationship with the same slope as the 0M urea data but a slightly higher rate. In fact, when all force data collected in this work are fit separately to eq. 2, the resulting x_U_^‡^ values are very close (Supplemental Table S1). Therefore, to minimize differences due to experimental error, these data were all fit together in a global analysis where x_U_^‡^ values were linked between experiments. The lines in Figures 2, 3 and 5b show the results of the global analysis (Supplemental Figure S1 compares the lines from individual fits and global fits). Table 1 reports the x_U_^‡^ value and the ln *k*_*U*_ value in the absence of urea and force for each construct resulting from the global fit. The x_U_^‡^ value from the global fit is within error of all x_U_^‡^ values from individual fits, validating our hypothesis that x_U_^‡^ is that same for all constructs studied here.

**Figure 3.**
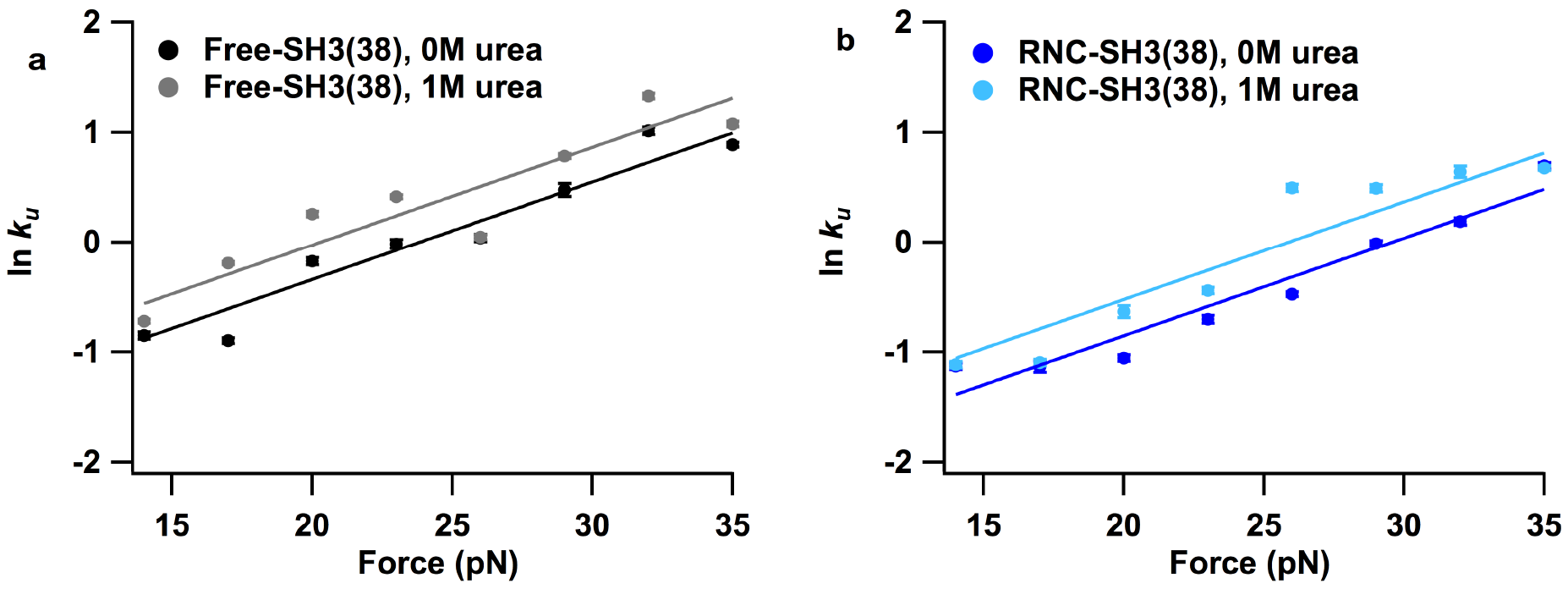
Chemo-mechanical unfolding analysis of free-SH3 and RNC-SH3. Plots of the natural log of the unfolding rate against force for (A) free protein with a 38 amino acid linker in 0 and 1M urea and for (B) RNCs with a 38 amino acid linker in 0 and 1M urea.

The effect of urea on unfolding rates is quantified by m_U_^‡^ values via equation 3:

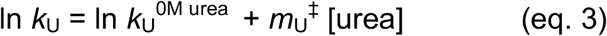

where [urea] is the molar concentration of urea and *k*_U_^0M urea^ is the unfolding rate in the absence of urea. The *m*_U_^‡^ value is related to the solvent accessible surface area exposed in unfolding to the transition state and so provides another parameter to compare transition states (*34, 35*). To determine *m*_U_^‡^ from chemo-mechanical unfolding data, we take the difference between ln *k*_U_^0pN^ for 1M urea and 0M urea data. The resulting *m*_U_^‡^ values for free-SH3(38) and RNC-SH3(38), reported in Table 1, are the same within error, indicating that the solvent accessible surface area exposed in unfolding to the transition state is the same for free-SH3 and RNC-SH3. This provides another parameter suggesting that the ribosome does not affect the folding transition state.

We have also determined *m*_U_^‡^ values for free-SH3 constructs in the absence of force from kinetic chevron experiments (Figure 1). Therefore, these m^‡^_U_ values can be used to compare the unfolding pathway in force experiments to the pathway in standard ensemble experiments. In fact, the *m*_U_^‡^ values for free-SH3(38) and RNC-SH3(38) force experiments are the same within error as the *m*_U_^‡^ values from chevron plots for all free-SH3 constructs used in this work, suggesting that the folding pathway in force experiments is the same folding pathway seen in standard ensemble experiments: SH3 folds through the same transition state on the ribosome that it folds through in bulk solution.

For example, in our previous work (*5*) different high force and low force pathways obtained when pulling in a shearing direction could be distinguished via their urea *m*_U_^‡^ values. On the other hand, pulling in an unzipping direction resulted in a similar *m*_U_^‡^ value to that obtained in the absence of force, suggesting a similar pathway (B/Z pathway). Thus, the similar *m*_U_^‡^ values obtained here from N- to C- pulling and bulk chevron experiments are consistent with SH3 unfolding through the same pathway in this pulling direction as the pathway seen in the absence of force (pathway B/Z) and that SH3 also folds through this pathway when tethered to the ribosome.

### Simulations of the effect of the ribosome on the folding of SH3 at different linker lengths

To gain further insight into the folding of src SH3 at different distances from the ribosome, we have determined the native state stability of the protein by molecular dynamics simulations of RNC-SH3 at different linker lengths. All the free energies are calculated under conditions where the stability of RNC-SH3 at long linker lengths (L=58aa) is equal to the stability of the isolated src SH3 domain (3.6 kcal/mol). Free energy profiles projected onto the fraction of native contacts (Figure 4 a and b), *Q* (*38*), show that src SH3 becomes more stable with longer linker lengths (Figure 4 a, b and c). The fraction of folded protein approaches ~100% (Figure 4c) after a sharp transition centered at L~34aa. In these simulations, at the linker length of L=38aa used in the experiments, the src SH3 domain has not completely emerged yet from the mouth of the tunnel. The domain is less stable (ΔΔG≅1.1 kcal/mol) than the wild type since several C-terminal residues are still in the tunnel. A representative set of folded states of SH3 from unbiased MD folding simulations are shown in Figure 4 d and e, from which we can observe that src SH3 is folding at the mouth of ribosome tunnel at linker length L=38aa and it folds completely outside of the tunnel at L=58aa. These simulations suggest that our experiments using a linker length of 38 amino acids is a good model to study the effect of folding close to the ribosome and it would be unproductive to shorten it further.

**Figure 4.**
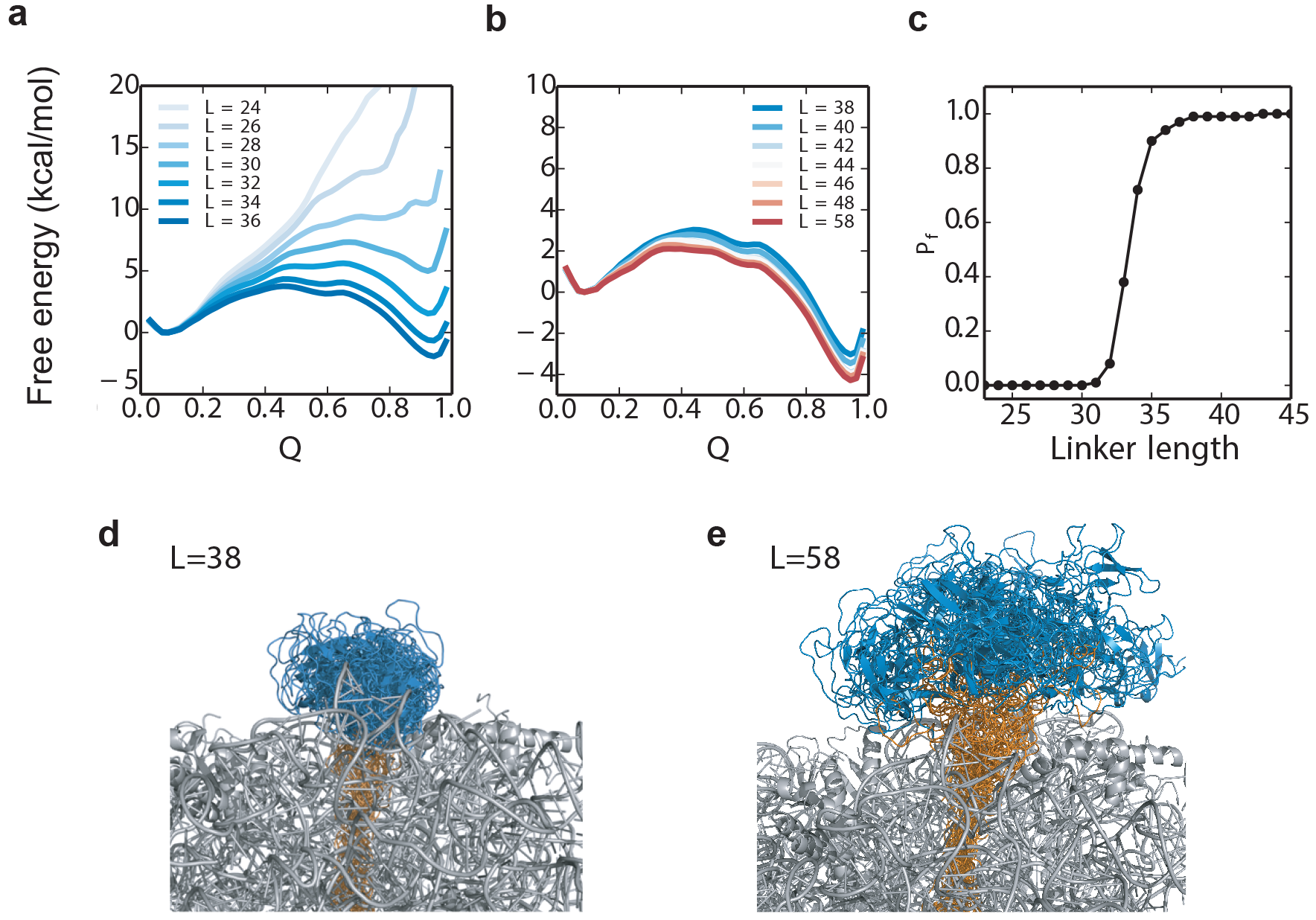
MD simulations of RNC-SH3 with different linker lengths. (A-B) Free energy profiles for src SH3 folding on the ribosome with different linker lengths (from 24 to 58 amino acids), Q is the fraction of native contacts. (C) Fraction of folded src SH3 population (P_f_) as a function of linker length. Ensemble of folded states of src SH3 with linker L=38aa (D) and L=58aa (E). Each ensemble contains conformations obtained from 2 microsecond MD simulations (saved every 40 nanoseconds) during which the protein remained folded.

### Characterizing folding pathways for free-SH3 and RNC-SH3 using point mutation

Like urea and force, the effect of mutation is related to the folding pathway (*39*). Therefore, a point mutation can be used to test our hypothesis that src SH3 unfolds through the same pathway on the ribosome and in bulk solution. In previous work, we determined the effect of various mutations on unfolding rates for the different folding pathways of SH3 and found that the L24A mutation significantly increases the unfolding rate for pathway B/Z (*5*). Here, we make the same L24A mutation on the free-SH3(38) and RNC-SH3(38) constructs to see if it also increases the unfolding rate in these experiments. We determine folding and unfolding rates for free-SH3(38) from a kinetic chevron plot fit to equation 1 (Figure 5a). Free-SH3(38) L24A behaves very similarly to src SH3 studied in previous work (without the tags and linkers) ‒ the mutation significantly destabilizes src SH3, primarily by increasing the unfolding rate.

**Figure 5.**
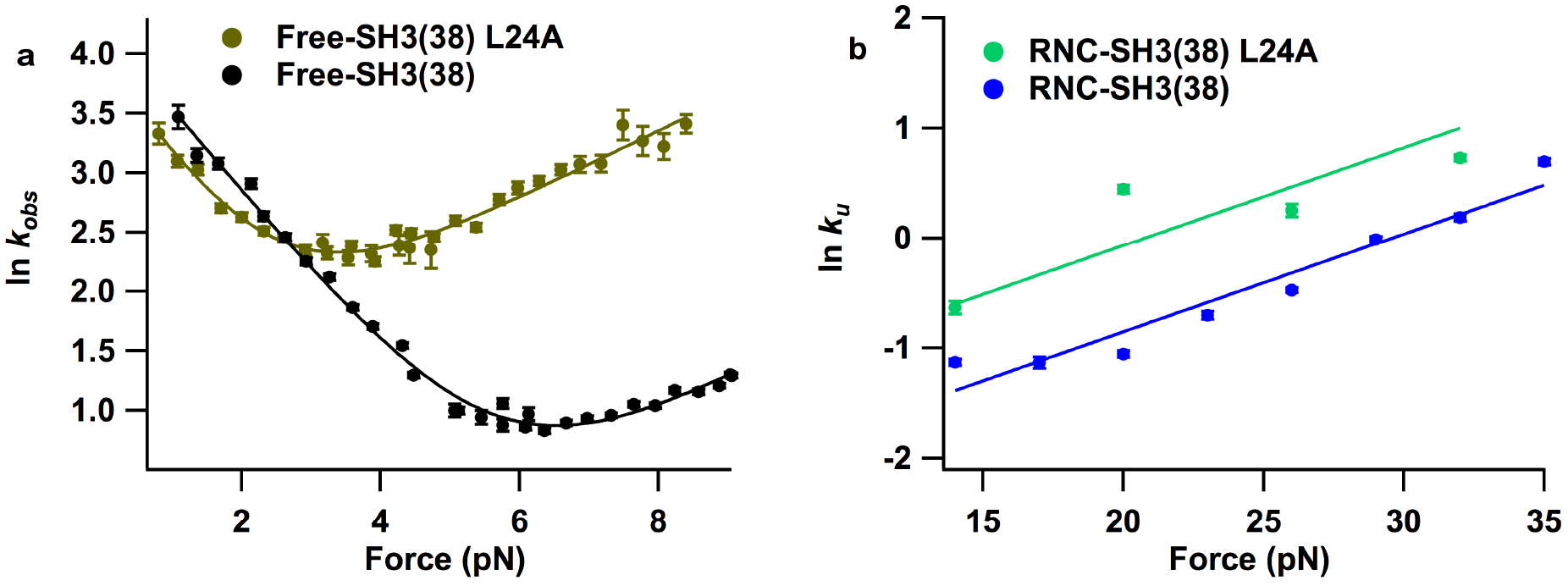
Effect of L24A mutation on unfolding rate of free-SH3 and RNC-SH3. (A) Kinetic chevron plots of natural log of observed relaxation rate against urea concentration for WT and L24A SH3 free protein construct. (B) Plots of the natural log of the unfolding rate against force for WT and L24A RNCs with a 38 amino acid linker.

For the RNC-SH3(38) L24A, we determine the unfolding rate as a function of force and find a linear relationship with a similar slope to the other constructs studied here (Figure 5b). Therefore, these data were included in the global analysis described above, linking all of the *x*^‡^_U_ values. Comparison to WT RNC-SH3(38) data shows that, as expected for pathway B/Z, the L24A mutation increases the unfolding rate for SH3 on the ribosome. Mutations are often used in a Φ-value analysis to characterize a transition state (*39*). This analysis requires determining the effect of mutation on the stability of the protein. Unfortunately, we are unable to determine the stability of src SH3 on the ribosome and so cannot calculate an experimental Φ-value. We are only able to measure unfolding rates for RNC-SH3 constructs and not folding rates because SH3 folds at very low forces where the optical tweezers have poor resolution. Instead, we can use simulations to determine Φ-values for src SH3 on and off the ribosome.

### Simulations of Φ-values for src SH3 on- and off- the ribosome

Previous works have shown that MD simulations with a coarse-grained model can reproduce the transition state ensemble of the isolated src SH3 domain (*40-43*). In our study, unbiased MD folding simulations at constant temperature are carried out for the isolated src SH3 domain as well as the RNC-SH3 constructs at different linker lengths, and the folding transition paths are used to estimate the Φ-values. We approximate Φ-values for each residue from folding transition paths, here defined as the segments of a trajectory between Q~0.3 and Q~0.6 (*44*). The simulated Φ-values are averaged from 50 transition paths for isolated src SH3, RNC-SH3(38) and RNC-SH3(58). The simulated Φ-value of residue 24 on the isolated src SH3 is 0.12 (Figure 6a), close to the experimental measurement (0.04) (*5*). Furthermore, the simulations reproduce well the overall F-value profile of src SH3 from previous experiments (Figure 6a) (*5, 45*). The largest discrepancy between simulation and experiments is at residue 44, which is also where the measurements obtained by two independent experimental groups differ the most. The Spearman rank correlation coefficient between the simulation and experimental data is 0.75 (Figure 6b), using Φ-values for mutations whose ΔΔG is larger than 7 kJ/mol (*46*). We also estimated the Φ-values for RNC-SH3(38) and RNC-SH3(58) (Figure 6c), obtaining Spearman correlations between these two cases and the experimental Φ values for the isolated src SH3 of 0.78 and 0.75 respectively. Taken together, these results suggest that the transition-state ensemble of src SH3 is not affected much by either the ribosome or the linker length.

**Figure 6.**
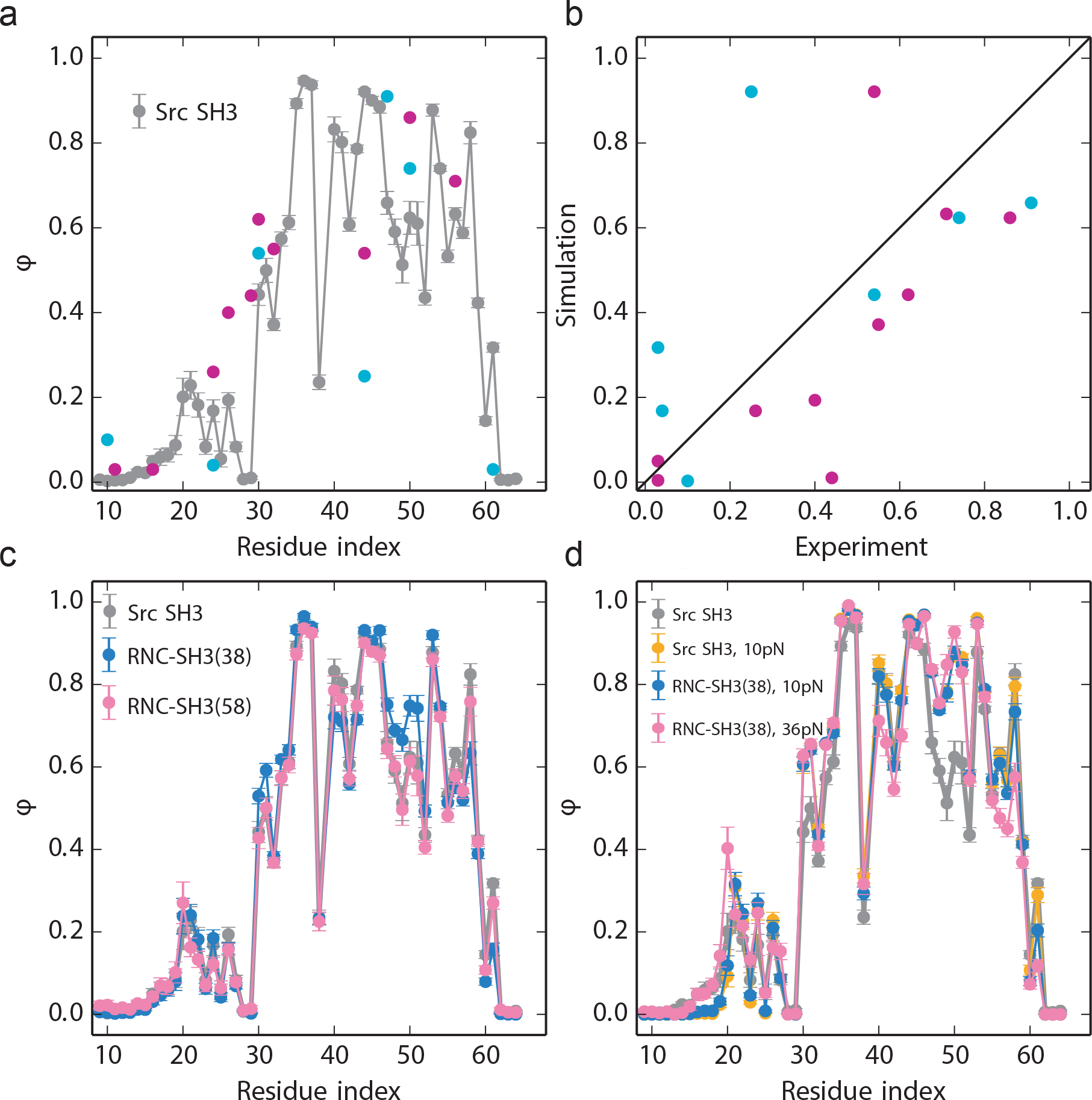
Calculated Φ values from MD simulations compared with experiments. A) Experimental Φ values of the isolated src SH3 domain from two experimental studies are shown in cyan (5) and purple (44) respectively. The simulated Φ values are calculated from the transition path ensemble of isolated src SH3 with no tags or linkers (gray). The correlation between simulated and experimental Φ values from A are shown in B. C) Simulated Φ values of RNC-SH3 with a 38 amino acid linker (blue) and RNC-SH3 with a 58 amino acid linker (red) compared with isolated src SH3 (gray). D) Φ values for the isolated src SH3 domain when it is subjected to 10pN pulling force are shown in yellow. Φ values for RNC-SH3 when the applied mechanical force is 10pN and 36pN are shown in blue and red respectively.

### Simulations of mechanical unfolding pathways on- and off- the ribosome

To make a direct comparison between simulations and the pulling experiments with free-SH3(38) and RNC-SH3(38), we estimated the force dependence of unfolding rates using MD simulations (Figure 7a). To be consistent with the experiments, in our simulations force is applied at the N and C- termini for both the isolated src SH3 domain and free-SH3(38). For RNC-SH3, force is applied to protein L17 of the ribosome and the N-terminus of the SH3 domain. As in our experiments, we find the same slope for the dependence of ln *k*_u_ on the pulling force for free-SH3(38) and RNC-SH3(38) (Figure 7b) (i.e. *x*^‡^_U_ is the same in the Bell model). This similarity supports the conclusion that the folding mechanism of src SH3 is the same on and off the ribosome. The value of *x*_U_^‡^ is small (~0.8 nm), suggesting a compact transition state. The value is slightly larger than experiment (~0.4 nm), which is most likely due to the coarse-grained model used. To determine whether the transition state ensemble of src SH3 changes under force, we compared the transition state ensemble (the Φ-values) of the isolated src SH3 with RNC-SH3 when force is applied (Figure 6d). The Φ-values obtained for isolated src SH3 at 10 pN are very similar to the Φ-values obtained for the RNC-SH3(38) at the same force (Figure 6d). Furthermore, Φ-values obtained for SH3-RNC (38) at larger force (36pN) do not differ significantly from the ones obtained at lower force (10pN, Figure 6d). Small differences are observed for residues 47-52.

**Figure 7.**
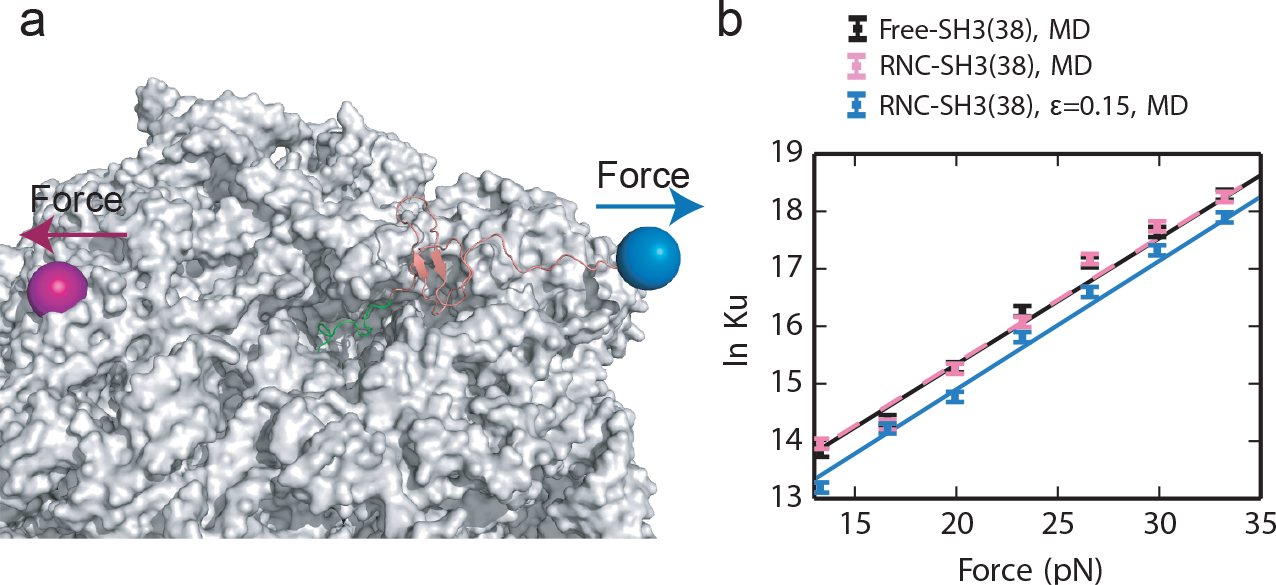
Mechanical unfolding of SH3 by MD simulations. A) Simulation setup for applying pulling force to RNC-SH3. SH3 is unfolded due to the mechanical force, which is applied at the N-terminus of SH3 (blue bead) and the C-terminus of protein L17 (red bead) of the ribosome (same as experiments). B) Force dependence of the unfolding rates for free SH3(L=38aa, black), RNC-SH3 (L=38aa, pink) and RNC-SH3 including attractive interactions between the nascent chain and ribosome (L=38aa, blue) from MD simulations. Black, pink and blue lines are fitted to the data of free SH3(L=38aa), RNC-SH3 (L=38aa) and RNC-SH3 (L=38aa, ε=0.15 kJ/mol) respectively.

So far, for the RNC constructs, we have made the simple approximation that the interactions between the src SH3 nascent chain and ribosome are given by short-range repulsion. The simulated mechanical unfolding rate (*k*_*U*_) for free-SH3(38) is the same as the RNC-SH3(38) (Figure 7b); however, the pulling experiments have demonstrated that the unfolding rate of src SH3 is slightly lower relative to free SH3(38) (Figure 2a), suggesting stabilization of the folded state relative to the unfolding transition state. We hypothesized that sufficiently weak attractive interactions between the ribosome and SH3 might stabilize the folded state and slow unfolding, similar to what has been observed in the case of protein folding in models of an attractive chaperonin cavity (*47*). We found that weak, non-specific attractive interactions can indeed slow down the unfolding of SH3 on the ribosome (Figure 7b). This may help to explain why the src SH3 unfolding rate is lower for the RNC than free SH3 (Figure 2a). Remarkably, the magnitude of the slowdown by ribosomal interactions obtained from simulation (Figure 7c) is very close to the slowdown in experiments (Figure 2a) where ln *k*_u_ decreases by 0.51 for RNCs.

## Conclusions

Using a variety of experimental and computational techniques, we have demonstrated that src SH3 folds via the same mechanism on the ribosome and free in solution. Experimentally, the effect of force, urea and point mutation on src SH3 unfolding kinetics are the same on and off the ribosome, strongly suggesting that the folding transition state is not altered. Simulations of src SH3 on and off the ribosome, both with and without force, support that the ribosome does not affect the folding pathway.

Together, these studies indicate that during translation, src SH3 does not fold until the entire domain sequence has reached the mouth of the ribosome exit tunnel. There are examples of domains that can partially fold before their entire sequence is exposed. Single molecule FRET experiments suggest that the α-helical N-terminal domain of HemK forms a compact, non-native state before the entire N-terminal domain sequence is outside of the exit tunnel (*16, 48*). Additionally, subunits of bacterial luciferase assemble co-translationally before the entire LuxB subunit is fully outside the exit tunnel, suggesting the interface required for assembly can fold independently of the C-terminus of luxB (*49*). Finally, some spectrin domains have been found to fold via a different pathway on the ribosome if partly folded N-terminal structures are stable enough to fold in isolation of the C-terminus (*17*). All of these proteins (HemK, luxB and spectrin domains) are composed primarily of α-helices, which are formed by short-range contacts. On the other hand, src SH3 is composed predominantly of β-strands, which require long-range contacts to form and so may prohibit folding before the entire domain sequence is outside the ribosome. This long-range contact order may limit the ability of the ribosome to alter the folding pathway via vectorial folding. Still, the ribosome can affect protein folding via chemical and coulombic interactions with the protein once it has emerged from the exit tunnel. These interactions are sufficient to alter the stability and kinetics of some stalled RNCs (*13, 14*) but, for src SH3, they are not sufficient to alter the folding transition state.

Recently, an arrest peptide based assay has been used to interrogate co-translational folding of another small β domain: the Ig domain, titin I27 (*19*). This assay involves monitoring the ability of constructs with different linker lengths to create sufficient force to overcome the SecM stalling sequence and continue translating. This force has been directly correlated with co-translational folding. These arrest peptide-based assays, together with cryo-electron microscopy, indicate that that I27 folds in the mouth of the ribosome. By simulating Φ-values for folding of I27 in the mouth of the ribosome they find a direct correlation to experimental Φ-values determined off the ribosome. Thus, in spite of the different experimental approaches, this work comes to a similar conclusion as we do ‒ the ribosome does not affect the folding pathway of their single Ig domain.

Therefore, the ribosome does not appear to play a significant role in determining the transition state for protein folding for the small src SH3 and I27 domains. Perhaps, the ribosome plays a more dominant role in altering protein folding pathways for larger multi-domain proteins. Due to the vectorial nature of folding, the N-terminal domains of a multi-domain protein can fold while the C-terminus is still sequestered in the ribosome. In firefly luciferase, the initial folding of the N-terminal domain has been shown to provide a scaffold that greatly speeds up the folding process of the native protein (*29*). Vectorial folding can also prevent aggregation-prone intermediates from forming, as in HaloTag, where an aggregation-prone intermediate observed in *in vitro* refolding experiments is not observed in co-translational folding experiments (*15*).

The fact that vectorial folding of protein domains can alter the folding pathway for multi-domain proteins highlights the importance of translation rates. The length of time that the N-terminal domains are outside of the ribosome exit tunnel before the C-terminus of the protein emerges determines how long the N-terminal domain has to fold in isolation of the C-terminus. Indeed, codon usage, which alters translation rates, has been found to affect protein folding efficiency and therefore downstream cellular processes (*23-25, 28*). If our observation that the ribosome does not greatly alter the rates of folding for small domains is general, then knowledge of the folding rates determined *in vitro* for individual protein domains together with codon translation rates can be used to assess whether or not an individual domain can fold before the protein is fully translated.

In sum, chemo-mechanical unfolding, together with molecular dynamics simulations of RNCs, have proven to be complementary techniques for probing folding pathways on the ribosome. Because folding on the ribosome is very complex, employing these techniques to study other proteins and other aspects of co-translational folding, like the presence of molecular chaperones, will help shed more light on this process.

## Materials and Methods

### Preparation of Free Protein Samples

Free SH3 constructs were expressed in *Escherichia coli* strain Rosetta/pLysS using the pET28a vector with the SH3 gene inserted using the NdeI and HindIII sites. This adds an N-terminal His Tag that was not removed in any of the RNC or free-protein constructs. The Avi tag was placed at the N-terminal end of this His tag so that it was at the N-terminus of the entire protein construct. All free SH3 constructs were purified using Ni-NTA resin followed by gel-filtration chromatography with a Hi-Load Superdex-75 16/60 column on a GE Healthcare AKTA purification system. Samples were stored in HKM buffer (25 mM HEPES-KOH pH 7.4, 150 mM KCl, 5 mM Mg-acetate). Free-protein samples to be used in optical trapping experiments were labeled at the N-terminal Avi tag with biotin and at the C-terminal ybbr tag with CoA cross-linked to a DNA oligo as described previously (*50*).

### Preparation of RNC samples

The ribosomes used to generate RNCs contain a C-terminal ybbr tag on protein L17. The *E. coli* strain producing these engineered ribosomes was a gift from Prof. Christian Kaiser (Johns Hopkins University). Engineered ribosomes were isolated from this strain as described previously (*51*). CoA cross-linked with a DNA oligo was attached to xthe ybbr tag on the ribosome as described previously (*13*). Linear DNA templates for *in vitro* transcription and translation (IVT) reactions containing the T7 promotor and RNC construct gene but lacking a stop codon were generated by PCR from plasmid DNA. To generate RNCs with biotin covalently attached to the N-terminal Avi tag, 50 μL IVT reactions were carried out using a ribosome-free IVT kit (PURExpress Δ Ribosome Kit, New England Biolabs) supplemented with 50 pmol ribosomes modified with the CoA-labeled DNA, 500 ng template DNA, 2 μL RNAse inhibitor, 10 mM MgCl2, 10 mM DTT, 2.5 mM ATP, 10 mM biotin and 62 μM birA. After incubation for 30 min at 37°C, IVT reactions were loaded onto a 125 μL 1M sucrose cushion in HKM and 2mM DTT (HKM+DTT) and centrifuged at 200,000 X g for 40 minutes at 4 °C. The ribosome pellets were washed three times with HKM+DTT and resuspended in 25 μL HKM+DTT. Aliquots were flash-frozen and stored at -80°C.

### Ensemble Kinetic Experiments

Kinetic data for the chevron plots were collected on a BioLogic SFM-400/MOS 200 stopped-flow fluorescence system. 150 μM unfolded protein (in 9M urea HKM buffer) or folded protein (in HKM buffer) was diluted 10-fold into final folding or unfolding conditions (HKM buffer, varying [urea]) at 25°C. Intrinsic tryptophan fluorescence signal was monitored by excitation at 295 nm and the emission was collected using a 360 nm bandpass filter.

### Optical Trap Experiments

On the day of experiments, RNC and free protein samples modified with the CoA-linked DNA oligo were ligated to a DNA oligo on a 2.1 μm carboxylic acid coated polystyrene beads as described previously(*13*) and these sample-beads were stored on ice. Immediately prior to experiments, anti-digoxigenin coated beads were incubated with a 2kbp biotin/digoxigenin coated DNA handle and streptavidin to create a DNA-handle bead as described previously(*30*).

Experiments were conducted using an optical tweezers instrument described in previous studies(*52, 53*). The optical trap is made of two coaxial, counter-propagating lasers holding the 3.2 μm anti-digoxigenin-coated bead at the focus. The 2.1 μm bead containing the RNC or free-protein samples was held by suction in a micropipette. Tethers were formed by bringing the beads in close proximity to each other to form a biotin-streptavidin linkage between the 2 kbp DNA handle and the biotinylated Avi tag. The micropipette is stationary and the trapped bead is manipulated by steering the optical trap, which samples data at 1 kHz and had a spring constant of ~0.08 pN/nm.

Unfolding rates for RNC and free-protein constructs were measured at a range of forces using force-jump experiments (*7*). For each force and urea condition, at least 6 different tethers were used to collect at least 70 force-jumps. Unfolding rates were determined by fitting the resulting dwell times to an exponential model as described previously (*6*).

### Molecular dynamics simulations

The ribosome stalls when the last Proline (of SecM) codon is positioned at the A site of the ribosomal peptidyl transferase center (*54*), therefore, in the simulation model, the last arrest-essential Proline of SecM is tethered to the last P atom in the A-site tRNA. The native interactions within SH3 domain are modelled using a coarse-grained structure-based model (*55*) based on the native structure of SH3 (pdbcode:1srl (*56*)), but with all of the contacts sharing the same contact energy. Each amino acid of SH3 or ribosome proteins is represented by one bead. Each RNA nucleotide of the ribosome is represented by three beads which are centered as the C4’, N3 and P atoms of the sugar, base and phosphate respectively. More details about the model have been described in a previous study (*19*). For the MD simulations, to obtain the free energy efficiently, umbrella sampling along reaction coordinate *Q* is applied, reweighting the results using WHAM (*57*). The Φ-values of SH3, RNC-SH3(38) and RNC-SH3(58) in Figure 6a are calculated based on the transition-path ensemble obtained by unbiased folding simulations at temperature 291K as previously described (*19, 44*). The Φ-values of SH3 and RNC-SH3(38) when SH3 is subject to mechanical forces (Figure 6d) are calculated based on transition-path ensemble obtained by unfolding simulations with mechanical force. All the MD simulations are carried out with a version of the software package Gromacs version 4.6.5 (*58*) modified to include the necessary 12-10-6 potential (*55*).

The attractive interactions between the nascent chain and ribosome are modeled by a Lennard-Jones potential:

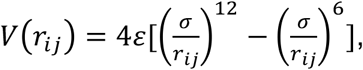

where *r*_*ij*_ is the distance between each bead pair *i* and *j. σ* is the range of interaction, which is set at 6 Å. *ε* is the strength of the interaction whose value is 0.15 kJ/mol.

## Acknowledgements

We thank Lisa Alexander, Daniel Goldman, Avi Samelson and Madeleine Jensen for guidance regarding *in vitro* transcription and translation reactions and DNA handle attachment. We thank Brendan Maguire for assistance with protein and ribosome purification and Jay Goodman for assistance with sample preparation. We would also like to thank the entire Marqusee lab for helpful discussions about this work. This research was supported by grants from the NSF (MCB 1616591) and the NIH (R01GM050945) to S.M. and an NIH fellowship to E.J.G. (F32GM110940). P.T. and R.B.B. were supported by the intramural research program of the National Institute of Diabetes and Digestive and Kidney Diseases of the National Institutes of Health. This study utilized the high-performance computational capabilities of the Biowulf Linux cluster at the National Institutes of Health, Bethesda, MD. (http://biowulf.nih.gov).

## References

1. A. Gershenson, L. M. Gierasch, Protein folding in the cell: challenges and progress. Curr Opin Struct Biol 21, 32–41 (2011).

2. M. Sarkar, A. E. Smith, G. J. Pielak, Impact of reconstituted cytosol on protein stability. Proc Natl Acad Sci U S A 110, 19342–19347 (2013).

3. I. Guzman, H. Gelman, J. Tai, M. Gruebele, The extracellular protein VlsE is destabilized inside cells. J Mol Biol 426, 11–20 (2014).

4. T. R. Sosnick, D. Barrick, The folding of single domain proteins--have we reached a consensus? Curr Opin Struct Biol 21, 12–24 (2011).

5. E. J. Guinn, B. Jagannathan, S. Marqusee, Single-molecule chemo-mechanical unfolding reveals multiple transition state barriers in a small single-domain protein. Nat Commun 6, 6861 (2015).

6. E. J. Guinn, S. Marqusee, Exploring the Denatured State Ensemble by Single-Molecule Chemo-Mechanical Unfolding: The Effect of Force, Temperature, and Urea. J Mol Biol 430, 450–464 (2018).

7. B. Jagannathan, S. Marqusee, Protein folding and unfolding under force. Biopolymers, (2013).

8. R. L. Baldwin, Temperature dependence of the hydrophobic interaction in protein folding. Proc Natl Acad Sci U S A 83, 8069–8072 (1986).

9. C. F. Wright, A. Steward, J. Clarke, Thermodynamic characterisation of two transition states along parallel protein folding pathways. J Mol Biol 338, 445–451 (2004).

10. E. P. O’Brien, B. R. Brooks, D. Thirumalai, Molecular origin of constant m-values, denatured state collapse, and residue-dependent transition midpoints in globular proteins. Biochemistry 48, 3743–3754 (2009).

11. L. D. Cabrita, C. M. Dobson, J. Christodoulou, Protein folding on the ribosome. Curr Opin Struct Biol 20, 33–45 (2010).

12. L. D. Cabrita et al., A structural ensemble of a ribosome-nascent chain complex during cotranslational protein folding. Nat Struct Mol Biol 23, 278–285 (2016).

13. C. M. Kaiser, D. H. Goldman, J. D. Chodera, I. Tinoco, C. Bustamante, The ribosome modulates nascent protein folding. Science 334, 1723–1727 (2011).

14. A. J. Samelson, M. K. Jensen, R. A. Soto, J. H. Cate, S. Marqusee, Quantitative determination of ribosome nascent chain stability. Proc Natl Acad Sci U S A 113, 13402–13407 (2016).

15. A. J. Samelson et al., Kinetic and structural comparison of a protein’s cotranslational folding and refolding pathways. Sci Adv 4, eaas9098 (2018).

16. W. Holtkamp et al., Cotranslational protein folding on the ribosome monitored in real time. Science 350, 1104–1107 (2015).

17. O. B. Nilsson et al., Cotranslational folding of spectrin domains via partially structured states. Nat Struct Mol Biol 24, 221–225 (2017).

18. O. B. Nilsson et al., Cotranslational Protein Folding inside the Ribosome Exit Tunnel. Cell Rep 12, 1533–1540 (2015).

19. P. Tian et al., The Folding Pathway of an Ig Domain is Conserved On and Off the Ribosome. bioRxiv, (2018).

20. A. Javed, J. Christodoulou, L. D. Cabrita, E. V. Orlova, The ribosome and its role in protein folding: looking through a magnifying glass. Acta Crystallogr D Struct Biol 73, 509–521 (2017).

21. A. M. Knight et al., Electrostatic effect of the ribosomal surface on nascent polypeptide dynamics. ACS Chem Biol 8, 1195–1204 (2013).

22. J. A. Farias-Rico, F. Ruud Selin, I. Myronidi, G. Von Heijne, Effects of protein size, thermodynamic stability, and net charge on cotranslational folding on the ribosome. bioRxiv, (2018).

23. A. K. Sharma, E. P. O’Brien, Non-equilibrium coupling of protein structure and function to translation-elongation kinetics. Curr Opin Struct Biol 49, 94–103 (2018).

24. G. N. Jacobson, P. L. Clark, Quality over quantity: optimizing co-translational protein folding with non-‘optimal’ synonymous codons. Curr Opin Struct Biol 38, 102–110 (2016).

24. G. Zhang, M. Hubalewska, Z. Ignatova, Transient ribosomal attenuation coordinates protein synthesis and co-translational folding. Nat Struct Mol Biol 16, 274–280 (2009).

26. W. M. Jacobs, E. I. Shakhnovich, Evidence of evolutionary selection for cotranslational folding. Proc Natl Acad Sci U S A 114, 11434–11439 (2017).

27. M. V. Rodnina, The ribosome in action: Tuning of translational efficiency and protein folding. Protein Sci 25, 1390–1406 (2016).

28. I. M. Sander, J. L. Chaney, P. L. Clark, Expanding Anfinsen’s principle: contributions of synonymous codon selection to rational protein design. J Am Chem Soc 136, 858–861 (2014).

29. J. Frydman, H. Erdjument-Bromage, P. Tempst, F. U. Hartl, Co-translational domain folding as the structural basis for the rapid de novo folding of firefly luciferase. Nat Struct Biol 6, 697–705 (1999).

30. D. H. Goldman et al., Mechanical force releases nascent chain-mediated ribosome arrest in vitro and in vivo. Science 348, 457–460 (2015).

31. J. Yin et al., Genetically encoded short peptide tag for versatile protein labeling by Sfp phosphopantetheinyl transferase. Proc Natl Acad Sci U S A 102, 15815–15820 (2005).

32. B. K. Kay, S. Thai, V. V. Volgina, High-throughput biotinylation of proteins. Methods Mol Biol 498, 185–196 (2009).

33. N. Ismail, R. Hedman, N. Schiller, G. von Heijne, A biphasic pulling force acts on transmembrane helices during translocon-mediated membrane integration. Nat Struct Mol Biol 19, 1018–1022 (2012).

34. E. J. Guinn, W. S. Kontur, O. V. Tsodikov, I. Shkel, M. T. Record, Probing the protein-folding mechanism using denaturant and temperature effects on rate constants. Proc Natl Acad Sci U S A 110, 16784–16789 (2013).

35. J. K. Myers, C. N. Pace, J. M. Scholtz, Denaturant m values and heat capacity changes: Relation to changes in accessible surface areas of protein unfolding. Protein Sci. 4, 2138–2148 (1995).

36. G. I. Bell, Models for the specific adhesion of cells to cells. Science 200, 618–627 (1978).

37. C. Bustamante, Y. R. Chemla, N. R. Forde, D. Izhaky, Mechanical processes in biochemistry. Annu Rev Biochem 73, 705–748 (2004).

38. R. B. Best, G. Hummer, W. A. Eaton, Native contacts determine protein folding mechanisms in atomistic simulations. Proc Natl Acad Sci U S A 110, 17874–17879 (2013).

39. A. R. Fersht, A. Matouschek, L. Serrano, The folding of an enzyme. I. Theory of protein engineering analysis of stability and pathway of protein folding. J Mol Biol 224, 771–782 (1992).

40. C. Clementi, H. Nymeyer, J. N. Onuchic, Topological and energetic factors: what determines the structural details of the transition state ensemble and “en-route” intermediates for protein folding? An investigation for small globular proteins. J Mol Biol 298, 937–953 (2000).

41. A. R. Lam et al., Parallel folding pathways in the SH3 domain protein. J Mol Biol 373, 1348–1360 (2007).

42. F. Ding, N. V. Dokholyan, S. V. Buldyrev, H. E. Stanley, E. I. Shakhnovich, Direct molecular dynamics observation of protein folding transition state ensemble. Biophys J 83, 3525–3532 (2002).

43. P. I. Zhuravlev, M. Hinczewski, S. Chakrabarti, S. Marqusee, D. Thirumalai, Force-dependent switch in protein unfolding pathways and transition-state movements. Proc Natl Acad Sci U S A 113, E715–724 (2016).

44. R. B. Best, G. Hummer, Microscopic interpretation of folding φ-values using the transition path ensemble. Proc Natl Acad Sci U S A 113, 3263–3268 (2016).

45. D. S. Riddle et al., Experiment and theory highlight role of native state topology in SH3 folding. Nat Struct Biol 6, 1016–1024 (1999).

46. I. E. Sánchez, T. Kiefhaber, Origin of unusual phi-values in protein folding: evidence against specific nucleation sites. J Mol Biol 334, 1077–1085 (2003).

47. D. Thirumalai, G. H. Lorimer, Chaperonin-mediated protein folding. Annu Rev Biophys Biomol Struct 30, 245–269 (2001).

48. E. Mercier, M. V. Rodnina, Co-translational Folding Trajectory of the HemK Helical Domain. Biochemistry, (2018).

49. Y. W. Shieh et al., Operon structure and cotranslational subunit association direct protein assembly in bacteria. Science 350, 678–680 (2015).

50. H. N. Motlagh, D. Toptygin, C. M. Kaiser, V. J. Hilser, Single-Molecule Chemo-Mechanical Spectroscopy Provides Structural Identity of Folding Intermediates. Biophys J 110, 1280–1290 (2016).

51. G. Spedding, Ribosomes and Protein Synthesis: A Practical Approach. G. Spedding, Ed., (Oxford Univ. Press, Oxford, 1990).

52. S. B. Smith, Y. Cui, C. Bustamante, Optical-trap force transducer that operates by direct measurement of light momentum. Methods Enzymol 361, 134–162 (2003).

53. C. Cecconi, E. A. Shank, C. Bustamante, S. Marqusee, Direct observation of the three-state folding of a single protein molecule. Science 309, 2057–2060 (2005).

54. H. Muto, H. Nakatogawa, K. Ito, Genetically encoded but nonpolypeptide prolyl-tRNA functions in the A site for SecM-mediated ribosomal stall. Mol Cell 22, 545–552 (2006).

55. J. Karanicolas, C. L. Brooks, The origins of asymmetry in the folding transition states of protein L and protein G. Protein Sci 11, 2351–2361 (2002).

56. H. Yu, M. K. Rosen, S. L. Schreiber, 1H and 15N assignments and secondary structure of the Src SH3 domain. FEBS Lett 324, 87–92 (1993).

57. S. Kumar, J. M. Rosenberg, D. Bouzida, R. H. Swendswn, P. A. Kollman, The weighted histogram analysis method for free-energy calculations on biomolecules. I. The method. Journal of Computational Chemistry 13, 1011–1021 (1992).

58. B. Hess, C. Kutzner, D. van der Spoel, E. Lindahl, GROMACS 4: Algorithms for Highly Efficient, Load-Balanced, and Scalable Molecular Simulation. J Chem Theory Comput 4, 435–447 (2008).

